# Distinct neural dynamics underlie the encoding of visual speed in stationary and running mice

**DOI:** 10.1101/2021.06.11.447904

**Authors:** Edward A B Horrocks, Aman B Saleem

## Abstract

Sensory experiences are often driven by an animal’s self-motion and locomotion is known to modulate neural responses in the mouse visual system. This modulation is hypothesised to improve the processing of behaviourally relevant visual inputs, which may change rapidly during locomotion. However, little is known about how locomotion modulates the temporal dynamics (time courses) of visually-evoked neural responses. Here, we analysed the temporal dynamics of single neuron and population responses to dot field stimuli moving at a range of visual speeds using the Visual Coding dataset from the Allen Institute for Brain Science^1^. Single neuron responses had diverse temporal dynamics that varied between stationary and running sessions. Increased dynamic range and more reliable responses in running sessions enabled faster, stronger and more persistent encoding of visual speed. Population activity reflected the temporal dynamics of single neuron responses, including their modulation by locomotor state - neural trajectories of population activity made more direct transitions between baseline and stimulus steady states in running sessions. The structure of population coding also changed with locomotor state – population activity prioritised the encoding of visual speed in running, but not stationary sessions. Our results reveal a profound influence of locomotion on the temporal dynamics of neural responses. We demonstrate that during locomotion, mouse visual areas prioritise the encoding of potentially fast-changing, behaviourally relevant visual features.

## Introduction

Sensory experiences are often driven by an animal’s self-motion. For example, an animal’s movement through an environment can drive rapid changes to visual inputs in the form of visual flow. The real-time encoding of visual motion is therefore critical for an animal to understand and control its interaction with an environment. Indeed, mice can accurately respond to moving visual stimuli at sub-second timescales^2^ and such rapid behavioural responses are likely to be of special ecological importance during active locomotion, where mice may be evading predators^3^ or hunting prey^4^.

Previous work in the mouse has demonstrated that locomotion modulates the neural encoding of visual features in various ways. For example, during locomotion neurons tend to have elevated firing rates^(5–7^, produce more reliable visual stimulus-evoked responses^8–10^ and exhibit stimulus-specific tuning gain^11^ and altered stimulus tuning preferences^12,13^. Locomotion is also associated with a reduction in pairwise spike-count correlations within mouse visual cortex^10,14,15^. Collectively, these results have been hypothesised to improve information processing capacity for behaviourally relevant visual inputs^15^. During locomotion these inputs may change rapidly, yet little is known about how locomotion influences the temporal dynamics (time courses) of visually-evoked neural responses. Consequently, how locomotion associated changes in temporal dynamics affect the real-time encoding of visual features, including visual speed, is not known. We therefore investigated the temporal dynamics of single neuron and population responses in mouse visual areas to large-field visual stimuli moving at a range of visual speeds.

We discovered distinct neural dynamics underlying the encoding of visual speed in stationary and running mice. Single neuron responses had diverse temporal profiles that varied between stationary and running sessions. Increased dynamic range and more reliable responses in running sessions enabled faster, stronger and more persistent encoding of visual speed. Population activity reflected the temporal dynamics of single neuron responses, including their modulation by locomotor state. Intriguingly, locomotor state also altered the structure of visual speed encoding by population activity – dominant factors of population activity prioritised the encoding of visual speed in running, but not stationary sessions.

## Results

To investigate the temporal response dynamics underlying visual speed encoding in stationary and running states we analysed the open-access Visual Coding dataset from the Allen Institute for Brain Science^1^. We focused our analyses on neural responses recorded while mice were presented dot field stimuli moving at one of seven visual speeds (0, 16, 32, 64, 128, 256 and 512°/s), in one of four directions (−45°, 0°, 45°, 90°), at 90% coherence (Figure 1a,b). Extracellular electrophysiology data was recorded using six Neuropixel probes^16^ targeting eight visual areas (Cortical areas: V1, LM, AL, RL, AM, PM and thalamic areas: dLGN and LP) of awake head-fixed mice (Figure 1c).

**Figure 1:**
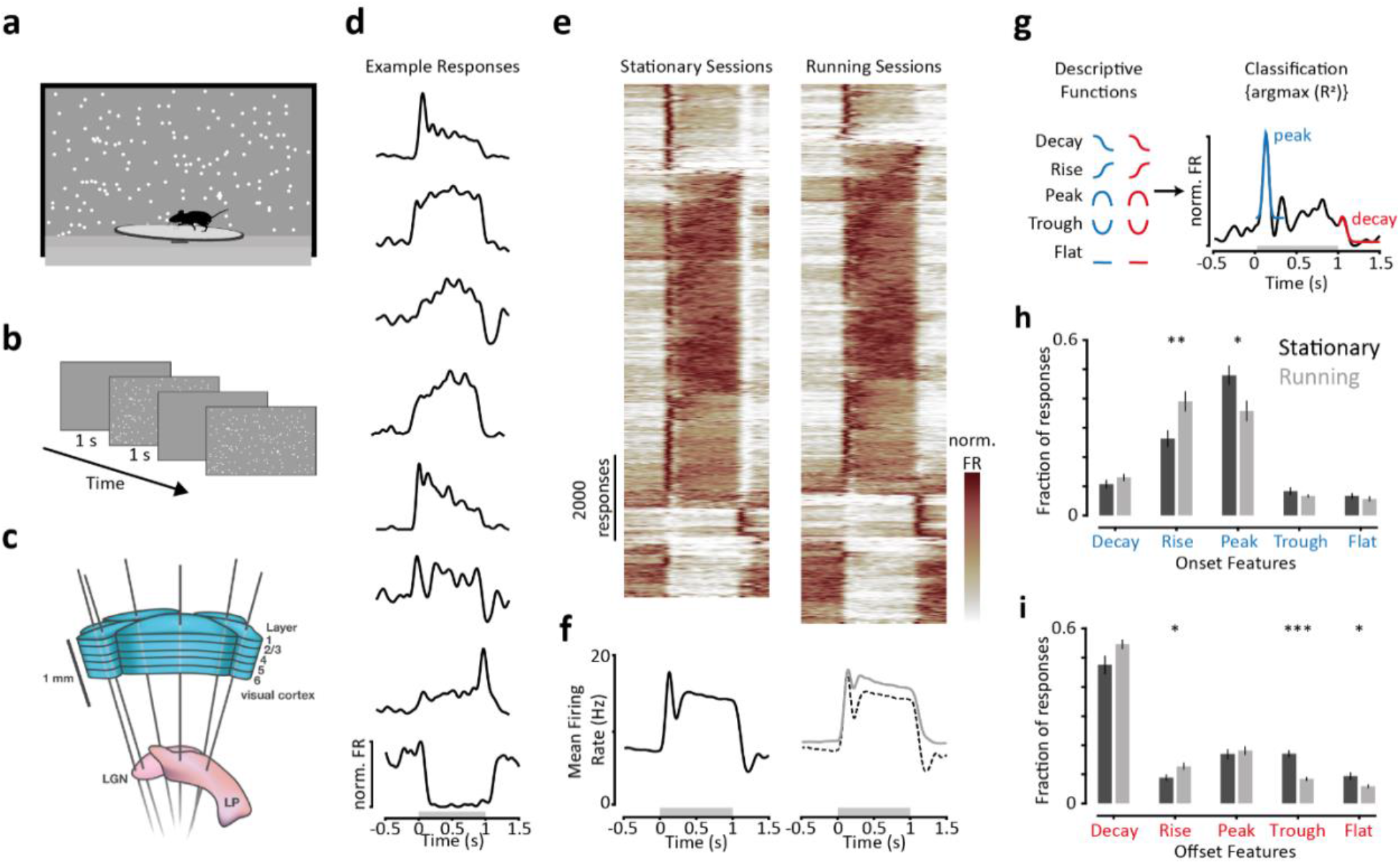
Neurons have distinct temporal response dynamics in running sessions. **a**. Illustration of experimental set-up. Moving dot fields were presented on a close-by monitor whilst mice were free to run on a disc. **b.** Moving dot field stimuli were presented for 1s and interleaved with a 1s mid-grey screen between trials. **c.** Six Neuropixel probes targeted eight visual areas (cortical: V1, LM, AL, RL, AM, PM; thalamic: dLGN, LP) simultaneously [Image from: https://portal.brain-map.org/explore/circuits/visual-coding-neuropixels]. **d.** Eight example peri-stimulus time histograms (PSTHs) from −0.5-1.5s. The stimulus was presented from 0-1s as indicated by the grey shaded region on the x-axis. **e.** All reliable PSTHs recorded in stationary (left panel) and running (right panel) sessions. Each row is an individual, normalised, reliable PSTH from a single neuron in response to a single stimulus condition (Stationary: n = 9,813/73,759 responses; running: n = 10,627/64,155 responses). Responses are sorted using a hierarchical sorting procedure. **f.** Mean reliable responses from stationary (left panel, black trace) and running (right panel, grey trace) sessions. The dashed line in the right panel is the mean stationary response, shown for comparison. **g.** Schematic of response feature classification. The onset (0-0.3s) and offset (1-1.5s) periods of each reliable response were classified as the bestfitting of five possible descriptive functions. An example response is shown along with the best-fitting functions overlaid. **h.** Mean fractions of onset features for all reliable responses recorded in stationary (black bars) and running (grey bars) sessions. Significance tested using Mann-Whitney U-test (n = 10 stationary, 7 running sessions). Error bars are ± sem. **i.** Same as **h** for offset features. *** p<0.001, ** p<0.01, * p<0.05.

During recordings mice were free to run on a disc (Figure 1a) and we compared the responses of single neurons and neural populations in stationary and running sessions. As mice in individual recording sessions tended to remain either stationary or running for the majority of the trials, we analysed only those trials recorded during the animal’s dominant locomotor state, discarding trials from the other locomotor state (Methods).

### Neurons have distinct temporal response dynamics in running sessions

Single-neuron responses had diverse temporal profiles which varied between stationary and running sessions. We characterised the temporal profiles of reliable responses based on the shapes of the peri-stimulus time histograms (PSTHs) of spiking activity (Figure 1d). We considered a response to be reliable if the PSTH had a similar temporal profile across multiple repeats of the same stimulus, quantified using cross-validation and a shuffle control (Methods). Reliable responses from both locomotor states could be excited, suppressed or a mixture of both; with large transients or more gradual changes in firing rate; and with peaks occurring throughout the response period (Figure 1d, e). The distributions of response shapes differed between stationary and running sessions (Figure 1e). For example, in running sessions there were fewer responses with strong transient onsets followed by lower sustained activity (top of Figure 1e), and more responses with sustained suppression (bottom of Figure 1e). Overall, the clearest differences in temporal response dynamics occurred immediately following stimulus onset (t = 0s) and offset (t = 1s). Responses recorded in stationary sessions had stronger transient increases of activity following stimulus onset and stronger suppression of activity following stimulus offset (Figure 1e, f).

The distributions of stimulus onset and offset response features differed between stationary and running sessions. We classified the onset (t = 0-0.35s) and offset (t = 1-1.5s) periods of each reliable response as the best-fitting of five possible descriptive functions (Decay, Rise, Peak, Trough and *Flat*; Figure 1g). Consistent with the mean responses (Figure 1f), there were significantly fewer responses with onset-peak (Figure 1h. Mean stationary = 47.9%, mean running = 35.7%. p = 0.04, n = 10 stationary, 7 running sessions) and offset-trough features (Figure 1i. Mean stationary = 17.1%, mean running = 8.5%. p < 0.001 n = 10 stationary, 7 running sessions) in running sessions. This difference was broadly consistent for responses evoked by different visual speeds (Supplementary Figure 1) and for neurons located in different visual areas (Supplementary Figure 2).

Having established that moving dot fields evoke distinct temporal response dynamics from neurons recorded in stationary and running sessions we next investigated whether there were any functional differences in the encoding of visual speed.

### Faster, stronger and more persistent encoding of visual speed by neurons in running sessions

Temporal profiles of visual speed tuning strength varied substantially across neurons (Figure 2a). To assess how the temporal dynamics of single neuron responses influenced functional encoding, we characterised visual speed tuning across different time windows (200ms sliding window; 10ms step size) using the cross-validated coefficient of determination (R^2^), which compares how well a tuning curve based on a training set of trials explains a tuning curve based on a held-out set of trials (Methods). Values greater than 0 indicate some degree of tuning for visual speed, and we classified tuning bouts as periods where tuning strength exceeded a threshold (R^2^≥0.1) for at least five consecutive times points (≥50ms step size). In many neurons tuning emerged quickly following stimulus onset and decayed quickly after stimulus offset (Figure 2a[i]). Other neurons exhibited transient tuning for visual speed, often during the stimulus onset (Figure 2a[ii]) or offset period (Figure 2a[iii]). Some neurons had multiple bouts of tuning (Figure 2a[iv]). Tuning for visual speed can therefore emerge throughout the response period, reflecting the diverse temporal dynamics present in single neuron responses.

**Figure 2:**
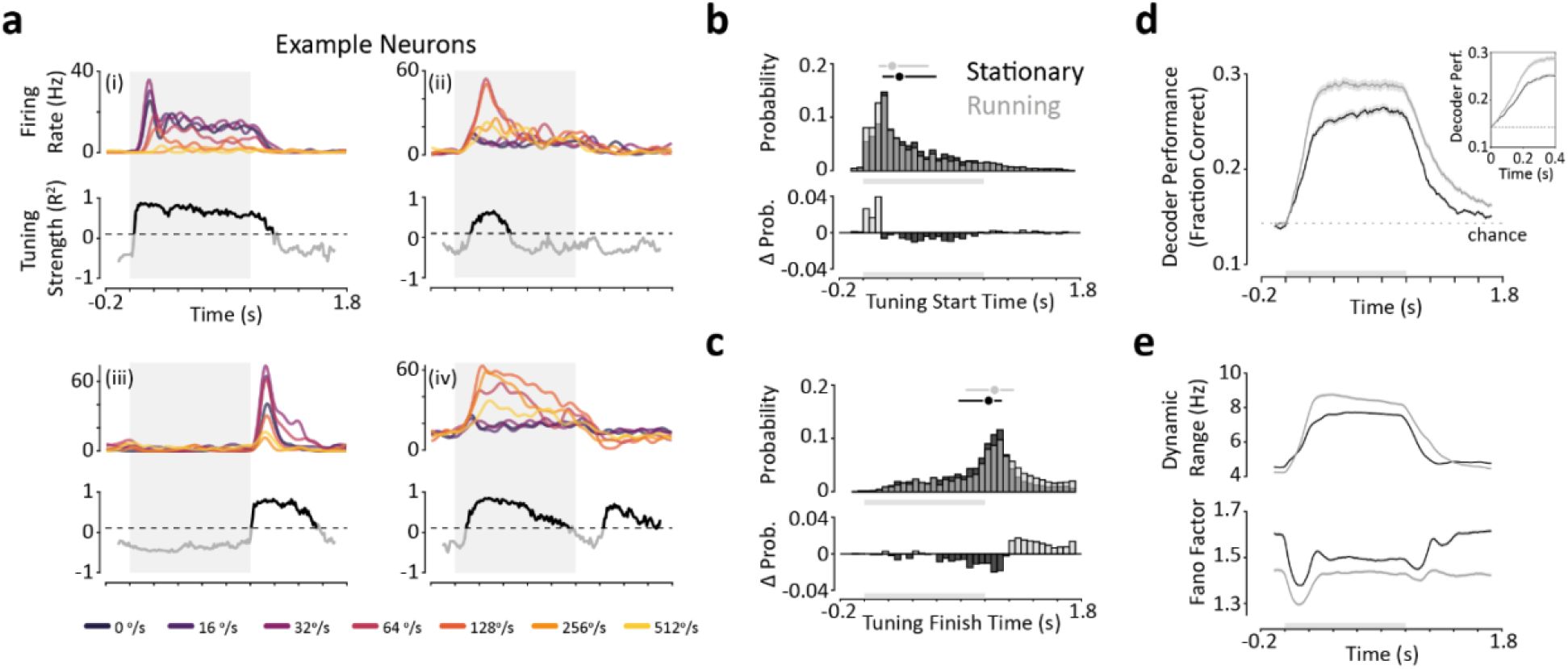
Faster and more persistent encoding of visual speed in running sessions. **a.** PSTH responses and tuning strength as a function of time for four example neurons (i-iv). *Top panels:* PSTHs for the seven visual speeds presented (in a single direction of motion). *Bottom panels:* vector of tuning strength calculated over a sliding window (200ms window size, 10ms step size). Dashed line represents the tuning threshold (R^2^=0.1) that separates tuned (black) and untuned (grey) periods of responses. Time on the x-axis corresponds to the window midpoint. Grey shaded region indicates stimulus presentation period. **b.** Tuning start times. *Top panel:* Probability histograms of tuning start times for neurons recorded in stationary (black) and running (grey) sessions. Circle and line plots above are median and interquartile ranges of respective distributions (Stationary: n = 5,080/10,537; running: n = 5,385/9,165 sets of responses with at least one tuning bout). Note that each neuron can contribute up to 4 sets of response depending on the number of valid directions of motion per session. *Bottom panel:* Difference histograms for the probability of neurons having a given tuning start time (positive values indicate a higher probability in running sessions). **c.** same as **b** for tuning finish times. **d.** Decoding performance (fraction correct responses) as a function of time for small populations of neurons (n = 10) recorded in stationary (black) and running (grey) sessions. Decoding was performed independently over time using a sliding window (200ms window size, 10ms step size). Time on the x-axis represents the window midpoint. Dashed line indicates chance level performance. Inset highlights the time period immediately following stimulus onset (0-0.4s). Shaded region indicates mean ± sem (Stationary: n = 306; running: n = 233 populations). **e.** Dynamic range (maximum firing rate - minimum firing rate) and mean Fano factor of single neuron responses as a function of time. Shaded region indicate mean ± sem for sets of responses recorded in stationary (black) and running (grey) sessions (Stationary: n = 10,537; running: n = 9,165 sets of responses).

Tuning for visual speed started earlier and finished later in neurons recorded during running. Tuning tended to start ~15% earlier during running sessions (Figure 2b; median tuning start time: stationary = 290ms, running = 240ms, Δ = −50ms. p=0.051; n = 10 stationary, 7 running sessions), a difference that reflected a higher fraction of neurons becoming tuned in the initial 150ms following stimulus onset. Tuning also finished significantly later during running (Figure 2c, median tuning finish time: stationary = 1010ms, running = 1080ms, Δ = +70ms. p = 0.006; n = 10 stationary, 7 running sessions), reflecting a tendency for neurons to remain tuned for more prolonged periods following stimulus offset. Overall a higher fraction of neurons were tuned for visual speed throughout the response period in running sessions, regardless of the specific tuning strength threshold used (Supplementary Figure 3). The distinct temporal response dynamics present in running sessions therefore underlie faster and more persistent tuning for visual speed.

Improved tuning enabled superior decoding of visual speed in running sessions (Figure 2d). We decoded visual speed from small populations of simultaneously recorded neurons (n=10) using a Poisson Independent Decoder^17^. The decoder assumes that neurons are independent and have Poisson firing rate statistics. The mean decoding performance of populations recorded in running sessions increased more rapidly following stimulus onset and decayed more slowly following stimulus offset (Figure 2d).

Decoding performance for visual speed also reached a higher steady state in running sessions, consistent with the larger fractions of neurons tuned during this period (Supplementary Figure 3). Thus, in running sessions information about visual speed emerged more rapidly following stimulus onset, reached a higher overall level and persisted for longer following stimulus offset.

**Figure 3:**
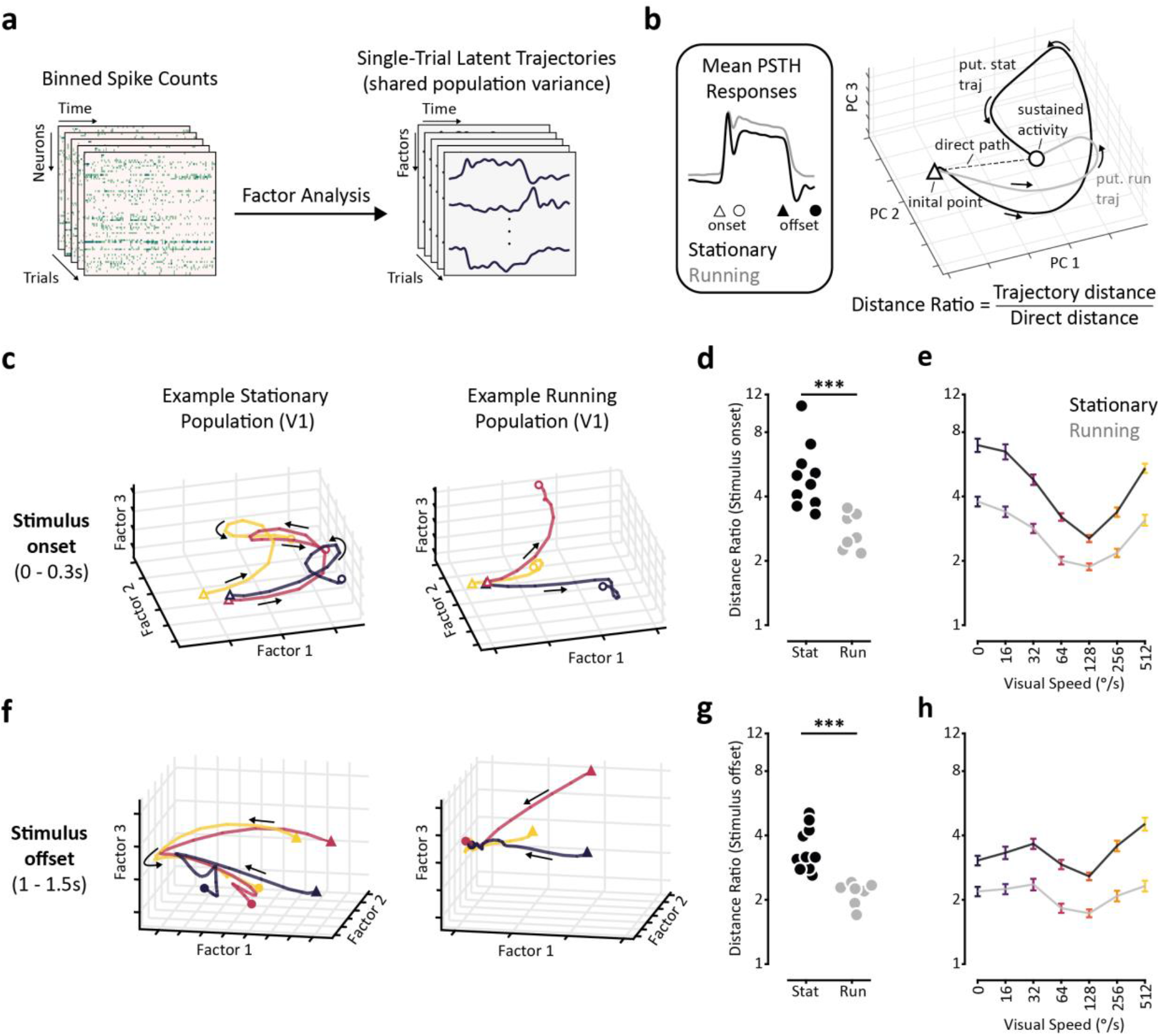
Population responses make more direct transitions between baseline and stimulus steady-states in running sessions. **a.** Illustration of Factor Analysis (FA). FA was performed on binned, square-root transformed and smoothed spike counts (left) to obtain latent factors of population activity (right). **b.** *Left panel:* mean single neuron reliable responses recorded in stationary and running sessions. *Right panel:* Hypothesis about population responses in stationary and running states - trajectories evolve from the initial point at 0s (indicated by the triangle) until reaching a steady-state of sustained activity (indicated by the circle). The direct path between the initial point and steady-state is shown by the dashed line. The putative stationary trajectory travels a greater distance before reaching the steady-state than the putative running trajectory, resulting in a higher Distance Ratio value for the putative stationary trajectory. *Bottom:* equation for Distance Ratio metric. **c.** Example trial-averaged latent trajectories of population activity from populations of V1 neurons recorded in a stationary (left) and running (right) session (only the top 3 factors are shown). Trajectories are shown for speeds 0°/s, 64°/s and 512°/s. **d.** Average Distance Ratios from stationary (black) and running (grey) sessions during the stimulus onset period. Significance tested using Mann-Whitney U-test (n = 10 stationary, 7 running sessions). **e.** Average distance ratios for the onset period as a function of visual speed for all stationary (black trace) and running (grey trace) populations (Stationary: n = 152; running: n =129). Error bars are ± sem. **f-h** Same as **c-e** for the offset period of responses. *** p<0.001.

We next asked whether the improved decoding and tuning for visual speed in running sessions was a result of increased dynamic range and/or response reliability. We assessed the separation of mean responses to different visual speeds (increased dynamic range) and the Fano factor (inverse of reliability) of neural responses. During running sessions the dynamic range of responses increased more rapidly following stimulus onset, reached a higher steady-state and decreased more slowly following stimulus offset (Figure 2e, top panel), similar to decoding performance (Figure 2d). The mean Fano factor of responses was also consistently lower throughout the response period for neurons recorded during running sessions (Figure 2e, bottom panel), reflecting more reliable stimulus-evoked responses. Interestingly, in running sessions mean Fano factor was lower in the baseline period before stimulus onset, indicating that spontaneous spiking activity was also more reliable, in agreement with previous work^8,9^. The improved encoding of visual speed by neurons in running sessions can therefore be attributed to a combination of increased dynamic range and more reliable responses.

The differences in visual speed tuning and decoding performance between stationary and running sessions varied between visual areas (Supplementary Figure 4). In areas V1, AM, PM and dLGN visual speed encoding was clearly enhanced in running sessions, as demonstrated by improved decoding performance. By contrast, in other visual areas decoding performance was often only subtly different between stationary and running sessions.

### Population responses make more direct transitions between baseline and stimulus steady-states in running sessions

Having explored the effects of running on single neuron responses we asked how these effects were reflected in population responses. We performed Factor Analysis on binned (20ms bin width) and smoothed (35ms Gaussian kernel) spike counts from populations of simultaneously recorded neurons in each visual area to obtain trajectories of population activity which evolved over time in latent factor space (Figure 3a). This approach produced latent factors of population activity with smooth time-varying single-trial trajectories. The extracted latent factors jointly represented the shared activity of neurons within a population, irrespective of visual speed. Overlapping groups of neurons contributed to individual latent factors with both positive and negative weights (Supplementary Figure 5). Individual latent factors thus represented both correlated and anti-correlated aspects of activity between groups of neurons. We initially focused our analyses on the trial-averaged temporal dynamics of population responses during the stimulus onset and offset periods, where we observed clear differences in single neuron responses between stationary and running sessions (Figure 1e, f).

We evaluated population responses based on the lengths of latent neural trajectories. We hypothesised that the increased prevalence of transient features (onset-peaks and offset-troughs) in single neuron responses recorded during stationary sessions would lead to more circuitous neural trajectories during the stimulus onset and offset period, when compared to running sessions. We compared the paths taken by neural trajectories of different populations of neurons using a distance ratio (Figure 3b). We define the Distance Ratio to be the distance travelled by a trajectory between the baseline and stimulus steady-states (or vice versa) divided by the most direct path between the two points (Methods). A distance ratio of 1 therefore represents a direct path, while higher ratios indicate less direct paths.

Neural trajectories of trial-averaged population activity took more direct paths between baseline and stimulus steady-states in running sessions (Figure 3c-h). We found that during the stimulus onset and offset periods of responses, trajectories of population activity from stationary sessions tended to take convoluted routes, often making reversals in direction and spiralling towards the eventual steady-state (Figure 3c, f). In contrast, neural trajectories from running sessions took more direct routes (Figure 3c, f). This was reflected in significantly lower Distance Ratios for trajectories obtained from running sessions during both the stimulus onset and offset periods (Figure 3d, g. Median onset distance ratio: Stationary = 4.79, Running = 2.57; p<0.001. Median offset distance ratio: Stationary = 3.17, Running = 2.25; p < 0.001. n = 10 stationary, 7 running sessions). This reduction in distance ratios was consistent across visual speeds (Figure 3e, h), but varied in magnitude between visual areas (Supplementary Figure 6).

### Latent factors of population activity robustly encode visual speed

We next investigated whether the more direct transitions between baseline and stimulus steady-states made by population responses in running sessions had functional correlates in the encoding of visual speed. We first assessed how well latent factors extracted using Factor Analysis encoded visual speed using Linear Discriminant Analysis (LDA) to decode visual speed. We compared decoding performance using binned spike counts of simultaneously recorded neurons and single-trial latent factor trajectories obtained using Factor Analysis, confining our analysis to the stimulus steady-state period of responses (0.5 - 1s post-stimulus onset, Methods). Decoder performance using latent factors was positively correlated with (rho = 0.80, p< 0.001, n = 282 populations from individual visual areas across 17 recording sessions) and superior to decoder performance using binned spike counts (Figure 4a. Signed-rank test: p<0.001). Latent factor spaces of population activity obtained using Factor Analysis therefore preserve information about visual speed present in binned spike counts.

**Figure 4:**
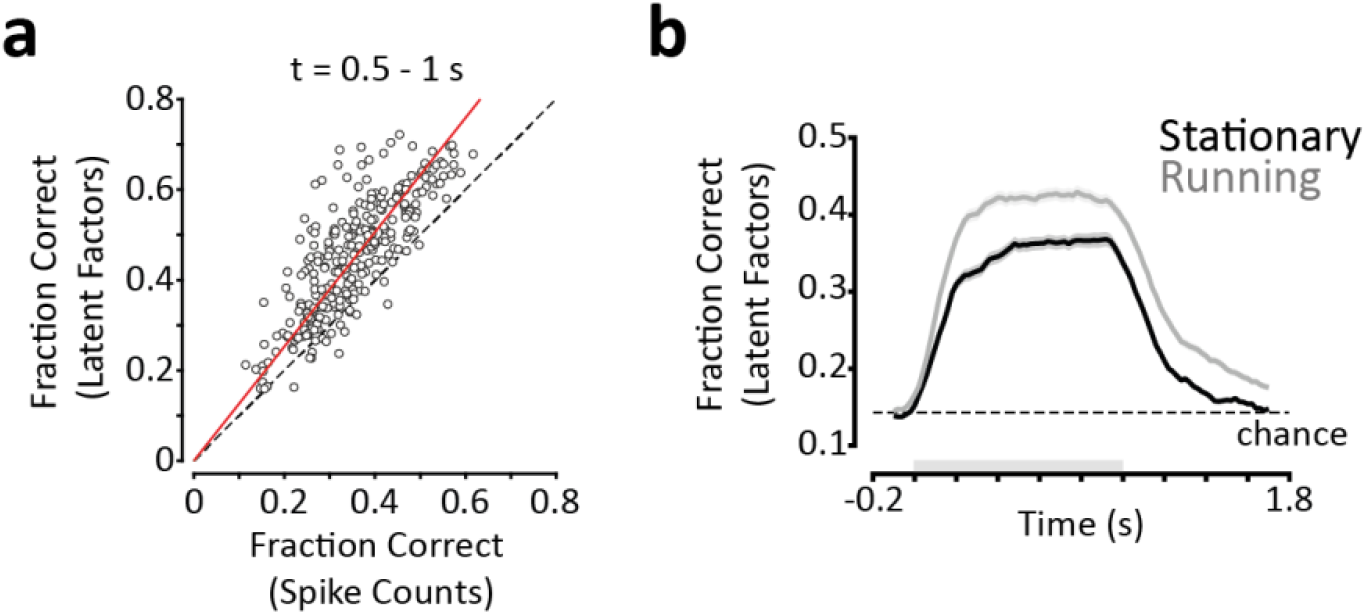
Latent factors of population activity robustly encodes visual speed. **a.** Scatter plot of LDA decoder performance (fraction correct predictions) for visual speed using smoothed binned spike counts (x-axis) and latent factor trajectories (y-axis). Each data point represents an individual population (n = 282). Overall performance was calculated as the mean decoder performance over multiple 200ms sliding windows (20ms step size) spanning 0.5-1s. **b.** Comparison of visual speed decoding performance using latent factors extracted from stationary and running sessions. Mean LDA decoding performance is plotted as a function of time for populations (Stationary: n = 225; running: n = 212) of fixed size (n=30 neurons) from stationary (black) and running (grey) sessions. Shaded regions are ± sem.

Visual speed could be decoded faster, to a higher maximum performance and for a more prolonged period using latent factors from running sessions (Figure 4b). We compared LDA decoding performance of latent factors obtained from stationary and running sessions (fixed population size of 30 neurons) across time. In running sessions, mean decoding performance increased more quickly following stimulus onset, reached a higher steady-state and decayed more slowly following stimulus offset (Figure 4b). The more direct transitions between baseline and stimulus steady-states made by population activity in running sessions therefore underlie faster and more persistent encoding of visual speed, similar to our results using a Poisson Independent Decoder and binned spike counts (Figure 2d).

### Population activity prioritises encoding visual speed during running

What aspects of population activity encode visual speed? Analyses of population activity using dimensionality reduction methods often focus on the dimensions which explain the largest amount of variance. However, it is not always apparent that these dimensions are the most relevant to the stimulus or behaviour of interest. Indeed, if the encoding of visual speed is prioritised during running then this may be reflected in the latent factors which encode it. We therefore assessed the relationship between the proportion of shared population variance explained by a latent factor (Figure 5a) and its ability to decode visual speed (Figure 5b), considering specifically whether this relationship varied between stationary and running sessions. To control for varying visual speed decoding performance by different populations of neurons we used a normalised performance metric to determine the relative performance of individual latent factors: the time-averaged decoding performance of each latent factor was divided by the time-averaged decoding performance of the full latent factor space extracted from the population of neurons (Figure 5b).

**Figure 5:**
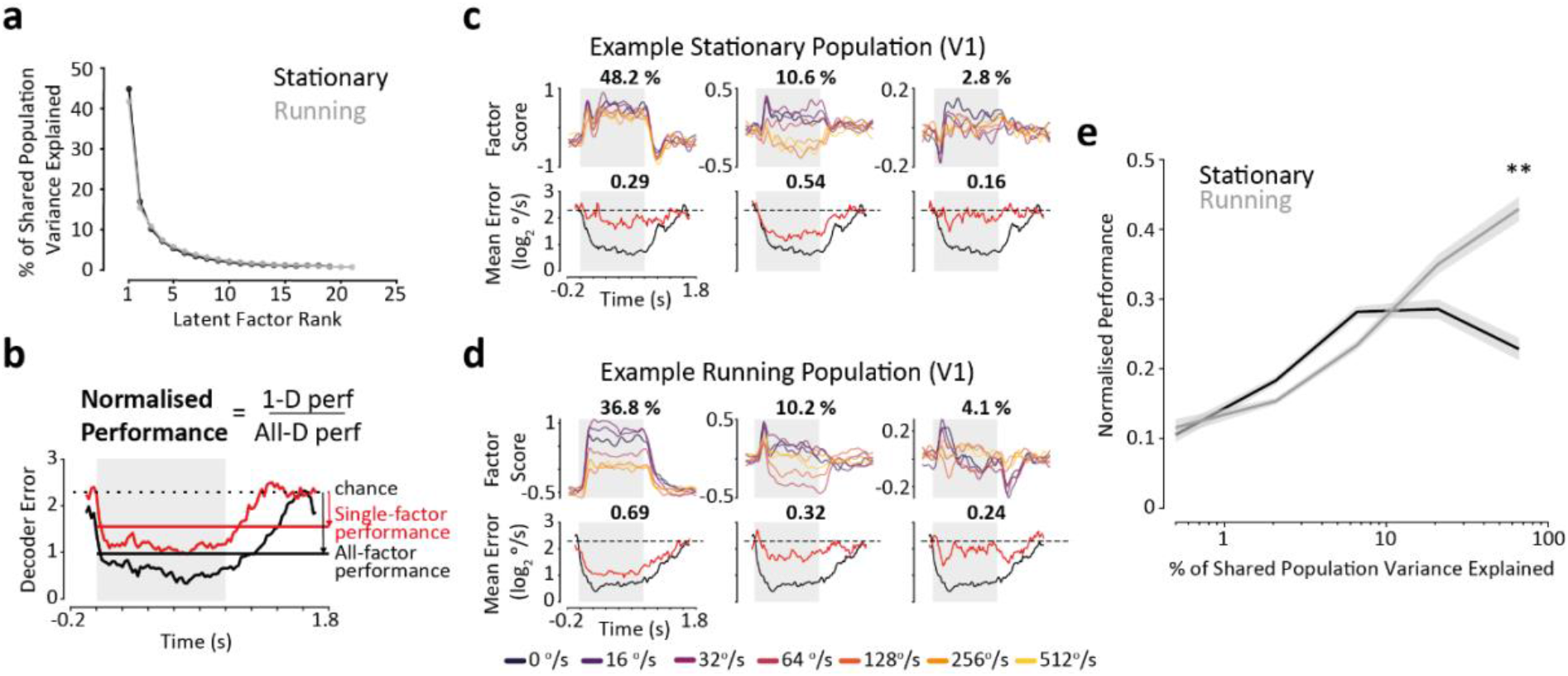
Population activity prioritises the encoding of visual speed during running. **a.** Mean percentage of shared population variance explained as a function of latent factor rank for populations of neurons recorded in stationary (black) and running (grey) sessions. **b.** Illustration of normalised performance metric. The time-averaged decoding performance of a single latent factor is divided by the time-averaged decoding performance of the full latent factor space extracted from a population of neurons. **c.** Example latent factors from a population of V1 neurons recorded in a stationary session. *Top row:* trial-averaged trajectories for each of the seven visual speeds presented. Numbers above each panel indicate the percentage of shared population variance explained by that factor. *Bottom row:* decoding performance as a function of time for the latent factor (red trace) and for the full latent factor space (black trace). Numbers above indicate the normalised performance of the latent factor. **d.** same as **c** for a population of V1 neurons recorded in a running session. **e.** Normalised performance as a function of percentage shared population variance explained (binned in log spaced intervals) for individual latent factors extracted from populations of neurons recorded in stationary (black) and running (grey) sessions (Stationary: n = 138; running: n = 128 populations). Shaded regions are mean ± sem, linearly interpolated between midpoints of percentage shared population variance explained bins. Statistical difference between stationary and running normalised performance values for individual bins tested using Mann-Whitney U-test (n = 10 stationary sessions, n = 7 running sessions). ** p < 0.01 (Bonferroni corrected). Grey shaded regions in b-d indicate the stimulus presentation period.

Population activity prioritised the encoding of visual speed during running. The relationship between the explained shared population variance and decoding performance of individual latent factors varied between stationary and running sessions (Figures 5c-e. Significant effect of explained variance on normalised decoding performance: F=30.173, df=4, p<0.001. Significant interaction between locomotor state and explained variance on normalised decoding performance: F=6.4192, df=4, p=0.002). In running sessions, decoding performance scaled monotonically with explained variance - the more variance a latent factor explained the better it was at decoding visual speed (Figure 5e). By contrast, in stationary sessions factors explaining intermediate proportions of variance performed best at decoding visual speed. These relationships are demonstrated by example ‘stationary’ and ‘running’ populations (Figures 5c, d). In the stationary population, whilst trajectories of the dominant latent factor clearly respond to the presence of a stimulus, the trajectories are not strongly dependent on visual speed and are therefore unable to decode visual speed effectively (Figure 5c). In contrast, the trajectories of the dominant latent factor in the example running population are clearly visual speed-dependent and are therefore able to effectively decode visual speed (Figure 5d). Indeed, normalised decoding performance of the dominant latent factors of population activity was significantly greater in running sessions (Figure 5e. p =0.002, Mann-Whitney U-test, n = 10 stationary, 7 running sessions). Thus, dominant factors of shared population activity prioritise the encoding of visual speed during running, but not stationary conditions. This difference was most pronounced in area V1 (Supplementary Figure 7).

## Discussion

Our results provide novel insights into how locomotion modulates the encoding of visual features by mouse visual areas. We demonstrate that during locomotion neurons exhibit distinct temporal dynamics in response to moving dot field stimuli. These distinct temporal dynamics have clear functional correlates, with increased dynamic range and response reliability during locomotion underlying faster, stronger and more persistent encoding of visual speed. We also demonstrate novel effects of locomotion on population activity. During locomotion population responses make more direct transitions between baseline and stimulus steady-states, reflecting the single neuron responses underlying them. Moreover, the structure of population coding for visual features depends on locomotor state - population activity prioritises the encoding of visual speed during running, but not stationary states. Our results provide further evidence that mouse visual areas adapt during locomotion to improve and prioritise the encoding of potentially fast-changing behaviourally relevant visual features, such as visual speed.

Our results demonstrate that visual speed can be decoded faster following stimulus onset during running, in agreement with previous work investigating the encoding of grating orientation and direction^15,18^. This speeding-up of encoding may enable mice to respond with faster reaction times to a rapidly changing visual environment during locomotion, where behavioural demands may be more immediate. We also found that visual speed can be decoded for longer following stimulus offset during running. This retention of information is reminiscent of persistent activity generally investigated in the context of working memory^19^, and may provide temporal context to changes in visual speed.

Our finding that locomotion alters the dynamics and structure of population coding for visual speed provides evidence that locomotor state has a profound impact on population coding for visual features in the mouse visual system. Indeed, dimensionality reduction methods are increasingly providing novel insights into the neural encoding of sensory features, including their modulation by behavioural state^20^. That visual speed encoding was prioritised by population activity during running raises the possibility that population activity adapts to behavioural priorities, such as the encoding of behaviourally relevant visual features. It has previously been suggested that locomotion induces a switch to more feed-forward processing in the mouse^21^, which would also suggest a coding prioritisation for external sensory variables. In contrast, during stationary periods population activity may prioritise more internal state variables, for example related to replay or planning^22^.

Our results demonstrate that the effects of locomotion on visual speed encoding vary between mouse visual areas (Supplementary Figure 4). We found substantial differences in the encoding of visual speed between mouse visual areas, including in the effects of locomotion. In areas V1, AM, PM and dLGN there were clear improvements in visual speed decoding during locomotion, whilst other areas showed only subtle differences between stationary and running sessions. The area-specific differences in decoder performance across time between locomotor states could generally be explained by area-specific changes in the dynamic range and mean Fano factor of responses. Different mouse visual areas are believed to be specialised for different functions owing to the varying response properties of neurons within them^12,23–25^. Indeed, the joint encoding of visual speed and running speed by neurons varies between visual areas^25,26^. Our results provide further evidence for the functional specialisation of mouse visual areas and moreover demonstrate their differential modulation by locomotor state.

What mechanisms are responsible for the distinct temporal response dynamics present in running sessions? Multiple neuromodulatory systems innervate mouse visual cortex and display increases in activity during locomotion^27^. These neuromodulatory systems have a range of effects during locomotion including altering the functional connectivity and intrinsic membrane properties of neurons^9,28–30^. Indeed, altered intrinsic membrane properties have been associated with more reliable spontaneous activity and stimulus-evoked responses^8,9^, which we also observed. Changes in functional connectivity have also previously been used to model changes in tuning properties during locomotion^31^. Thus, changes in neuromodulatory inputs to mouse visual areas, along with other top-down inputs, may underlie the distinct temporal dynamics we observed during locomotion.

In summary, we find profound effects of locomotion on the temporal dynamics of responses to visual stimuli. Our results highlight the importance of considering temporal aspects of neural encoding, especially in the context of active behaviours where timing can be crucial. We also provide new insights into population coding of visual features. Using dimensionality reduction we show that both the temporal dynamics and structure of population activity change during locomotion in order to prioritise and improve the encoding of potentially fast-changing, behaviourally relevant features such as visual speed.

## Acknowledgements

The authors declare no conflict of interest. This work was supported by The Sir Henry Dale Fellowship from the Wellcome Trust and Royal Society (200501); the Human Frontier in Science Program (RGY0076/2018), Biotechnology and Biological Sciences Research Council grant (R004765) to A.B.S.; and Biotechnology and Biological Sciences Research Council studentship to E.H.. We thank Saskia de Vries and Josh Siegle for discussions, and Sylvia Shroeder and Gordon Berman for comments on the manuscript.

## Author Contributions

This work was conceptualised by E.H. and A.B.S., Methodology, Software and Formal Analysis were by E.H., Visualization and Writing by E.H. and A.B.S, and Supervision and Funding Acquisition by A.B.S.

## Methods

### Experimental Data

We analysed in vivo extracellular electrophysiology recordings of mouse visual areas performed by the Allen Institute for Brain Science^1^. These recordings simultaneously targeted eight mouse visual areas (Cortical areas: primary visual cortex (V1), lateromedial cortex (LM), anterolateral cortex (AL), anteromedial cortex (AM), posteromedial cortex (PM). Thalamic areas dorsal lateral geniculate nucleus (dLGN), lateral posterior nucleus (LP)) using six Neuropixel probes^16^.

We analysed sessions where moving dot field stimuli were presented (*Functional Connectivity* stimulus-set). Dot fields consisted of ~200 3° diameter white dots moving across a mean-luminance grey background. In a given trial all the dots moved at one of seven visual speeds (0, 16, 32, 64, 128, 256, 512°/s) and in one of four directions (−45°, 0°, 45°, 90°; 0° is nasal to temporal motion, positive changes indicate clockwise rotation) at 90% coherence. Stimuli were repeated 15 times in a random order.

For each session and direction of motion we assessed whether at least 10/15 trials for every visual speed were viewed whilst mice were in a single locomotor state (stationary or running). We defined a trial as stationary if the mean running speed was <3cm/s during stimulus presentation and as running if the mean running speed was >5cm/s. Where sufficient trials were available for a given locomotor state, we analysed those trials and discarded trials from the other locomotor state in the session. Using this criteria we analysed 17/24 sessions: 10 sessions with stationary trials (mean number of directions of motion = 3.2) and 7 sessions with running trials (mean number of directions of motion = 3.6). There was no cross-over between sessions with sufficient stationary and running trials.

### Peri-Stimulus Time Histograms (PSTHs)

To obtain trial-averaged response profiles as a function of time (temporal response dynamics) we constructed peri-stimulus time histograms (PSTHs) for a given neuron and stimulus condition. We first calculated spike counts in 10ms time bins from 0.5s pre-stimulus onset to 0.5s post-stimulus offset (2s total). We then averaged the responses across trials and smoothed the resultant binned average response using a Gaussian kernel with 35ms standard deviation.

### PSTH reliability

We used cross-validation to determine the reliability of PSTHs across trials. The reliability metric determines whether the temporal profile of a PSTH using half of the available trials is significantly more similar to a PSTH using the other half of available trials compared to a PSTH constructed with shuffled spike times. Significantly reliable responses had fluctuations of firing rate which were consistent across trials whilst unreliable responses tended to have firing rate fluctuations that differed across trials.

For each cross-validation iteration, we randomly split trials into two equal sets and constructed two PSTHs (from 0-1.5s). We then calculated the time-summed Euclidean distance between the firing rates (spikes/s) of the two PSTHs, using Dynamic Time Warping^32,33^ with a maximum sample value of 2 bins (20ms). This was achieved using the Matlab function *dtw*. For each cross-validation iteration, we also randomly shuffled the spike times in one set of trials and constructed a PSTH as usual from these shuffled spike times. We then calculated the same distance measure between the normally constructed PSTH (from one half split of trials) and the shuffled-PSTH (from the other half split of trials). For each cross-validation iteration, we repeated this shuffling procedure 20 times.

After 50 cross-validation iterations we obtained a distribution of 50 normal distance measures and 1000 shuffled distance measures. We calculated the mean and standard deviation of the shuffled distance measures distribution to calculate a z-score for the mean of the actual distance measures distribution. Finally, we determined a PSTH response to be significantly reliable if this z-score value was ≤-1.645 (equivalent to p≤0.05).

### PSTH sorting

We sorted reliable PSTH responses using a hierarchical clustering procedure^34^. This sorting was performed on all reliable responses together. First, we normalised the firing rates of PSTHs to between 0 and 1. We then computed a dissimilarity matrix X_NxN_ (N = number of reliable responses) where X_i,j_ is the distance between PSTHi and PSTHj. As a distance metric we used Dynamic Time Warping (DTW) with a maximum sample value of 10 time bins (100ms) and the time bin-summed Euclidean distance. DTW enables the alignment of time series with similar shapes which are otherwise not precisely aligned in time. This approach was particularly important since we were comparing responses from multiple different visual areas with different onset latencies.

We used the dissimilarity matrix to perform hierarchical clustering using the unweighted average distance (Matlab function *linkage*). We then obtained the optimal ordering of PSTHs using an algorithm that minimises the sum of pairwise distances between neighbouring leaves of a dendrogram^35^ (Matlab function *optimalleaforder*).

### PSTH onset and offset feature classification

In order to classify the onset and offset features of PSTHs we first normalised the firing rates of PSTHs to between 0 and 1 and extracted the onset period (0-0.3s, where 0s was stimulus onset) and offset period (1-1.5s, where 1s was stimulus offset) of each normalised PSTH. We then separately fit a series of descriptive functions (*Decay*, *Rise*, *Peak*, *Trough* and *Flat*) to the onset and offset periods of each PSTH using the Matlab function *lsqcurvefit*.

Descriptive function were Gaussian of the form:

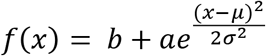

where *b* is a baseline firing rate parameter, *a* is an amplitude parameter, *μ* is the mean of the distribution and *σ* the standard deviation. *x* represents the zero-based time bins the functions were fit over.

Different parameter bounds were used to differentiate the descriptive functions. *Decay* and *Rise* features were described by two functions each (with positive and negative amplitudes). *Decay*, *Rise*, *Peak* and *Trough* features had appropriately bounded means (*μ*). The upper bound for the standard deviation (*σ*) was lower for *Peak* and *Trough* features to ensure well defined maxima or minima. *Trough* was differentiated from *Peak* by enforcing a negative amplitude (*a*). *Flat* was primarily defined by enforcing a small amplitude parameter (*a*).

Additionally, *Peak* and *Trough* features required the signed peak of the fitted function to have a prominence ≥0.2 (calculated using the Matlab function *findpeaks*). Moreover if the range of the normalised response over the onset or offset period was <0.2 (i.e. only a small amount of firing rate modulation) the feature was classified as *Flat* by default.

To compare the proportions of individual onset and offset features in stationary and running sessions we calculated the proportion of responses with a given response feature for each session and then performed the Mann-Whitney U-test on these values.

### Visual speed tuning strength and timing

We assessed the tuning strength of individual neurons over time using the cross-validated Coefficient of Determination (R^2^). For each 200ms sliding window (10ms step size) we performed 3-fold cross-validation where ⅔ of trials were sampled as a training set and the remaining ⅓ as a test set. For each iteration of cross-validation we constructed two models using the training set: a mean spike-count tuning curve model (*trained model*) based on the mean spike counts in response to each visual speed and a uniform *null model* calculated as the mean spike count to all visual speeds combined. Using the test set we constructed a mean spike-count tuning curve model (*test model*) in the same way as the *trained model*. We then determined how well the *trained model* and *null model* could predict the *test model* by calculating the sum-of-squared residuals between them. The coefficient of determination was then calculated with the following equation:

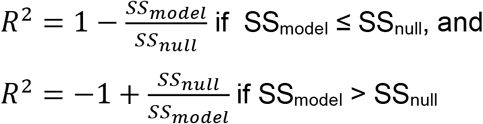

where SSmodel is the sum of squared residuals between the *trained model* and the *test model* and SSnull is the sum of squared residuals between the *null model* and the *test model*.

We then computed the mean R^2^ value over the 3 cross-validations, using a unique set of test trials on each iteration. We repeated this process 10 times with different random splits of train and test trials to obtain 10 estimates of R^2^. We calculated the mean of these 10 values as the final estimate of R^2^.

We considered a neuron to be tuned over a given time interval if it’s R^2^ value ≥0.1 for at least 5 consecutive sliding windows. Tuning start times were taken as the midpoint of the first sliding window of the first valid tuning interval. Tuning finish times were taken as the midpoint of the final sliding window of the final valid tuning interval.

To compare the tuning start and finish times from stationary and running sessions we found the median tuning start and finish time from each session and performed the Mann-Whitney U-test on these values.

#### Dynamic Range

The dynamic range of a set of responses was calculated as the minimum firing rate subtracted from the maximum firing rate across responses to all visual speeds for a given time window.

#### Fano Factor

Fano factor was calculated as 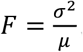, where σ^2^ and μ are the variance and mean of a set of spike counts, respectively. Mean Fano factor was calculated by averaging the Fano factor of responses across visual speeds for a given time window.

### Poisson Independent Decoder (PID)

To decode visual speed from spike counts of small populations of neurons we used a Poisson Independent Decoder^17^ (PID). For each session we generated multiple populations of neurons by sampling 10 neurons at a time without replacement from individual visual areas. We decoded visual speed in each direction of motion independently.

For each population we binned the spike times of the neurons into 10ms bins (−0.2−1.8s) to obtain binned spike counts for each neuron and trial. In order to compare the decoding performance of multiple populations we downsampled the number of trials used for decoding to 10 per stimulus condition (10 trials x 7 speeds = 70 trials in total). We performed decoding independently across time using a sliding window (200ms window size, 10ms step size). In each window we used leave-one-out cross-validation where visual speed-labelled spike counts from all but one trial per stimulus condition were used to train the decoder (9 trials x 7 speeds = 63 trials) and the spike-counts of the remaining trials (1 trial x 7 speeds = 7 trials) were used to test the decoder. This process was repeated 10 times until the decoder had predicted the visual speed of all 70 trials. Performance in each window was calculated as the fraction of correctly predicted trials. For each population we averaged performance over the available directions of motion.

For each trial visual speed was predicted by maximising the following log likelihood distribution:

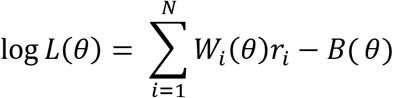

Where: *i* indexes over *N* neurons, *W_i_*(*θ*) is the logarithm of the mean spike count of neuron *i* for a given visual speed *θ* (learned from a training set of trials), *r_i_* is the number of spikes produced by neuron *i* during the trial being predicted and B(*θ*) is a bias term calculated as the sum of the mean spike counts for a given visual speed *θ* in the population (i.e. a correction for heterogeneous coverage of visual speeds by the tuning curves of a population).

### Factor Analysis

We obtained smooth, single-trial trajectories of population activity using Factor Analysis (FA). We considered populations with >30 valid neurons, where a neuron was considered valid if it had a mean firing rate >1Hz. We performed FA on responses to the seven visual speeds jointly, but to each direction of motion independently.

FA is defined as:

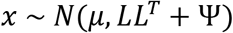

Where: *x* (*n* x 1) is vector of spike counts from *n* neurons; *μ* (*n* x 1) is a set of mean spike counts from the same *n* neurons; *L* (*m* x n) is the loading matrix which maps the *m*-dimensional latent variable to the spike counts of *n* neurons and Ψ(*n* x *n*) is a diagonal matrix of independent neuron variance. We estimated *μ, L* and Ψ using expectation-maximisation^36^ with code modified from the DataHigh Matlab toolbox^37^. FA explicitly separates shared covariance of the spike counts of a population of simultaneously recorded neurons from the independent spike count variance of individual neurons^38,39^. The low dimensional latent factor space obtained using FA therefore represents the shared population variance of a population of simultaneously recorded neurons.

We first binned the spike times of each neuron into 20ms bins (−0.2s−1.8s; 100 bins total). We then square-root transformed and smoothed the resultant vectors using a Gaussian kernel with a 35ms standard deviation^38^. We then used 3-fold cross-validation to find the dimensionality *m* of the latent variable which maximised the likelihood of the data^39^. Once we had obtained a value of *m* we defined *m*_opt_ as the smallest number of dimensions required to explain 95% of the shared population variance in the *m*-dimensional model. We then obtained a final latent factor space for a population of neurons by performing Factor Analysis with an *m*_opt_-dimensional latent variable.

### Distance ratios of trial-averaged trajectories

We analysed the paths taken by trial-averaged latent neural trajectories using an *m*opt-dimensional weighted Euclidean space. Each dimension of the space was a latent factor and its corresponding weight was the proportion of shared population variance explained by the factor. We defined the initial point of onset trajectories as the coordinate in this space at 0s (i.e. the first time bin following stimulus onset). The stimulus steady-state was defined as the cloud of points obtained from the trajectory during the interval 0.5-1s. We calculated the distances of individual points in the cloud to the centre of mass of the cloud and defined the maximum distance as SS_max_. We consider a trajectory to have reached steady-state if its distance to the centre of mass of the steady-state was less than SS_max_ for at least 10 consecutive time bins (i.e. 200ms total).

Offset trajectories were analysed in the same way with the exception that the initial point of the trajectory was defined as the coordinate in latent factor space at 1s (i.e. the first time bin following stimulus offset) and the baseline steady-state was defined using the cloud of points obtained from the trajectory between 1.5s-1.8s.

Distance Ratios were calculated as the cumulative distance travelled by a trajectory between the initial point and point of entry into the steady-state divided by the shortest Euclidean distance between the same two points:

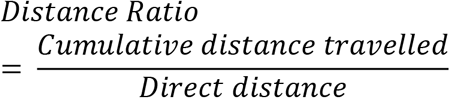

To compare the distance ratios obtained from stationary and running sessions we calculated the mean Distance Ratio (separately for stimulus onset and offset periods) across all populations recorded within a session (i.e. from different visual areas and in response to different directions of motion) and performed the Mann-Whitney U-test on these values.

### Decoding using Linear Discriminant Analysis

We used Linear Discriminant Analysis (LDA) to decode visual speed using the single trial trajectories of latent factor spaces, single trial trajectories of individual latent factors and binned and smoothed spike counts. Decoding was performed independently across time using a 200ms sliding window (20ms step size). When decoding using latent factors we took the mean values of single trial trajectories in a given time window as the predictor variables. When decoding using smoothed spike counts we simply calculated the mean binned spike-counts in the window and used those as predictor variables. In each sliding window we performed leave-one-out cross-validation where all but one trial for each visual speed were used to train the LDA model (Matlab function *fitdiscr*). The trained model then made predictions about the visual speed (Matlab function *predict*) of the test trials (1 trial x 7 speeds = 7 trials). This process was repeated until each trial had been predicted by the decoder.

To compare LDA decoding performance of visual speed using single-trial latent trajectories and smoothed spike counts (i.e. the input to FA) we performed a paired decoding analysis. For each population of neurons we decoded visual speed separately using latent trajectories and smoothed spike counts as predictor variables. For each decoding analysis we found the mean decoding performance (fraction correctly predicted trials) over the time window 0.5 – 1 s (i.e. the stimulus steady-state). We used Pearson’s correlation and the Wilcoxon signed rank test to compare performance using the two predictor variables.

To compare LDA decoding performance of visual speed using single-trial latent trajectories obtained from populations of neurons recorded in stationary and running sessions we performed FA on a fixed number of neurons at a time (N = 30), sampling without replacement. We also downsampled the number of trials to perform decoding on to 10 trials per stimulus condition (10 trials x 7 visual speeds = 70 trials total). We used the fraction of correctly predicted trials to assess decoder performance.

### Normalised performance of latent factors

To assess the decoding performance of individual latent factors relative to the full latent factor space obtained from a population of neurons we performed LDA decoding as usual using 1-dimensional latent trajectories (individual latent factor) and *m*opt-dimensional latent trajectories (full latent factor space) to get the time-varying decoding performance. Here we used geometric mean decoding error (log2[actual/predicted]) to obtain a more precise measure of decoding performance. For simplicity we enforced log2(0) = 3 so that the visual speeds presented were equally spaced in log2 space. In order to normalise the decoding performance of individual latent factors to the full latent factor space we first calculated a time-averaged decoding error (from 0-1.8s) for both of these predictor variables:

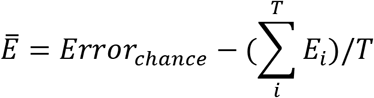

Where: *Error_chance_* is the average expected error from random predictions, *E_i_* is the mean decoder error during time window *i* and T is the total number of time windows.

The normalised performance of an individual latent factor was then calculated as:

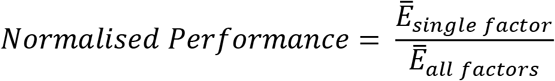

Where the subscript *single factor* denotes an individual latent factor and *all factors* the full latent factor space. Since some populations showed poor overall performance decoding visual speed we excluded populations with *Ē_all factors_* > 2 from further analysis.

To compare the relationship between % explained shared population variance and normalised performance of individual latent factors in stationary and running sessions we performed a repeated-measures ANOVA, with normalised performance as a response variable, % explained shared population variance as a within-subjects factor and locomotor state as a binary categorical between-subjects factor. For each population we found the mean normalised performance of latent factors binned according to their explained shared variance (5 logarithmically spaced out bins with edges = [0, 1, 3.16, 10, 31.62, 100] %). For each session we then calculated the median normalised performance for each of these 5 bins. The repeated-measures ANOVA was then performed using these 5 values for each session\ as the response variables. P-values were corrected using the Greenhouse-Geisser correction.

## Supplementary Figures

**Supplementary Figure 1.**
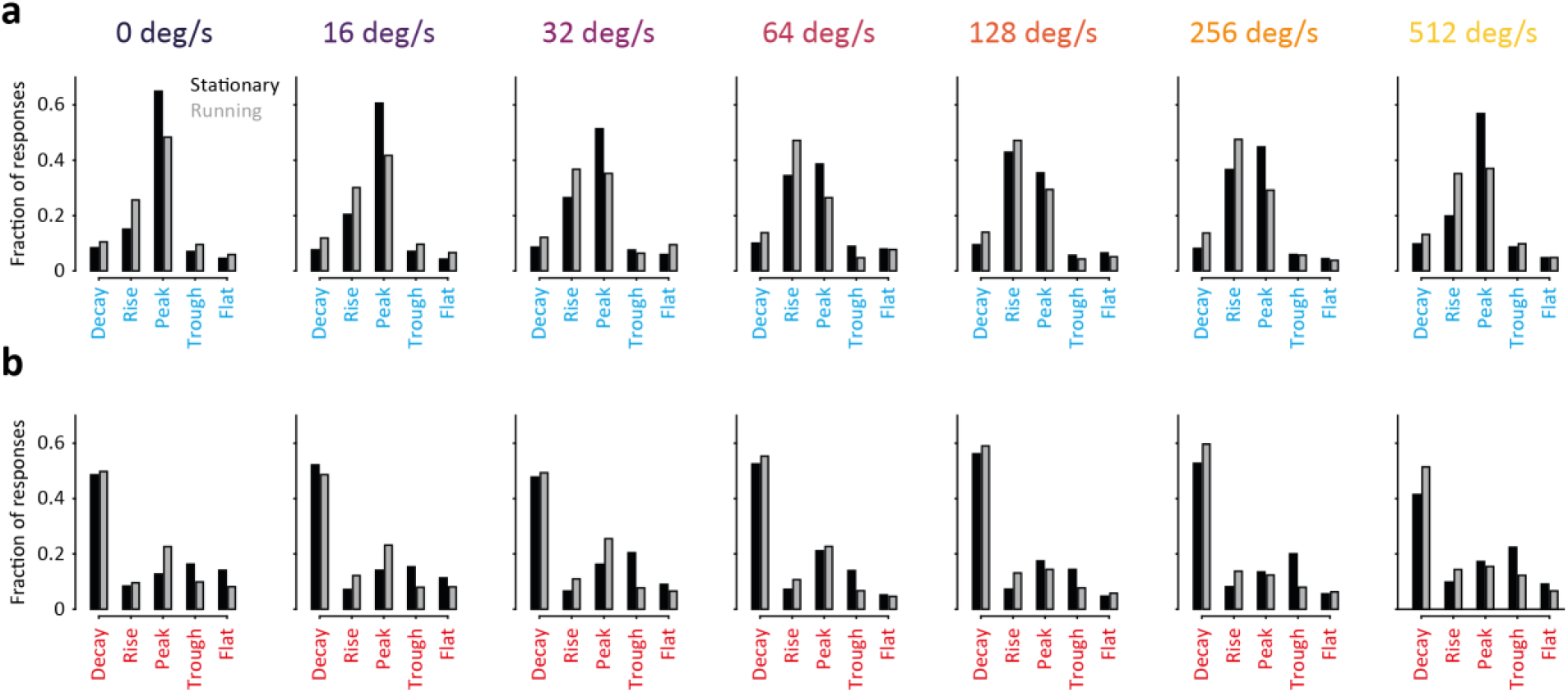
Differences in temporal response dynamics between locomotor states for each visual speed. **a.** Related to **Figure 1h** for individual visual speeds. Each panel shows the distributions of onset response features for all reliable responses evoked by each visual speed, pooled across stationary (black bars) and running (grey bars) sessions (Number of reliable responses [stationary sessions / running sessions]: 0°/s [1,220 / 1,301], 16°/s [1,254 / 1,479], 32°/s [1,423 / 1,675], 64°/s [1,974 / 2,099], 128°/s [1,872 / 1,811], 256°/s [1,191 / 1,281], 512°/s [906, 981]). The title above each plot indicates the visual speed. **b.** Same as **a** for offset response features. Related to **Figure 1i** for individual visual speeds.

**Supplementary Figure 2.**
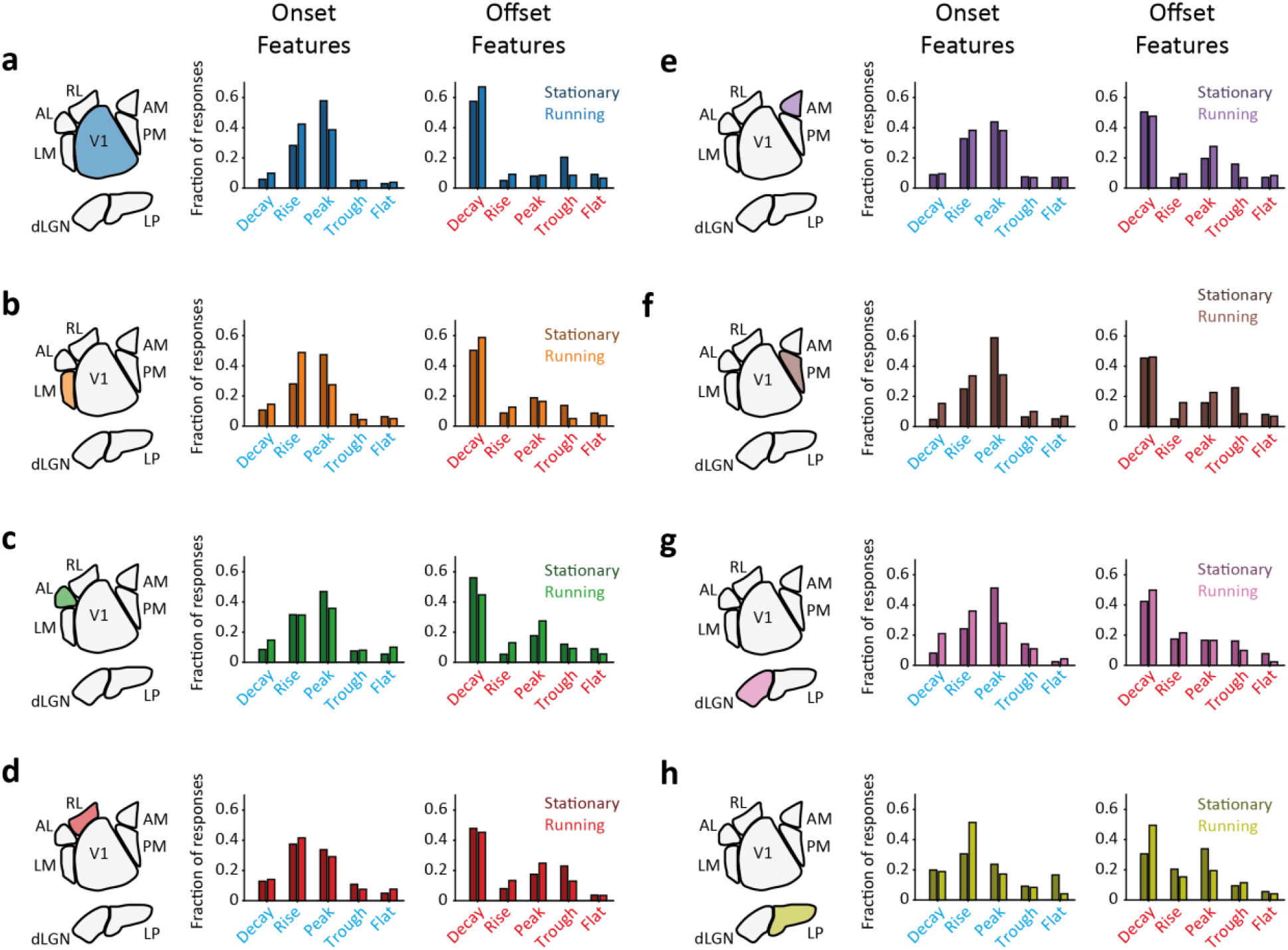
Differences in temporal response dynamics between locomotor states for each visual area. **a-h.** Related to **Figure 1h, i** for individual visual areas. Proportions of onset and offset response features in all reliable responses recorded in each visual area (Number of reliable responses [stationary sessions / running sessions]: V1 [2,415 / 2,879], LM [2,752 / 1,532], AL [1,212 / 2,343], RL [441 / 592], AM [1,268 / 1,641], PM [974 / 984], dLGN [185 / 390], LP [566 / 266]). *Left panel:* illustration of visual area anatomical location relative to other visual areas. The relevant visual area is colour-filled *Right panel*: proportions of different onset and offset response features for all reliable responses recorded in stationary (dark shaded bars) and running (light shaded bars) sessions.

**Supplementary Figure 3.**
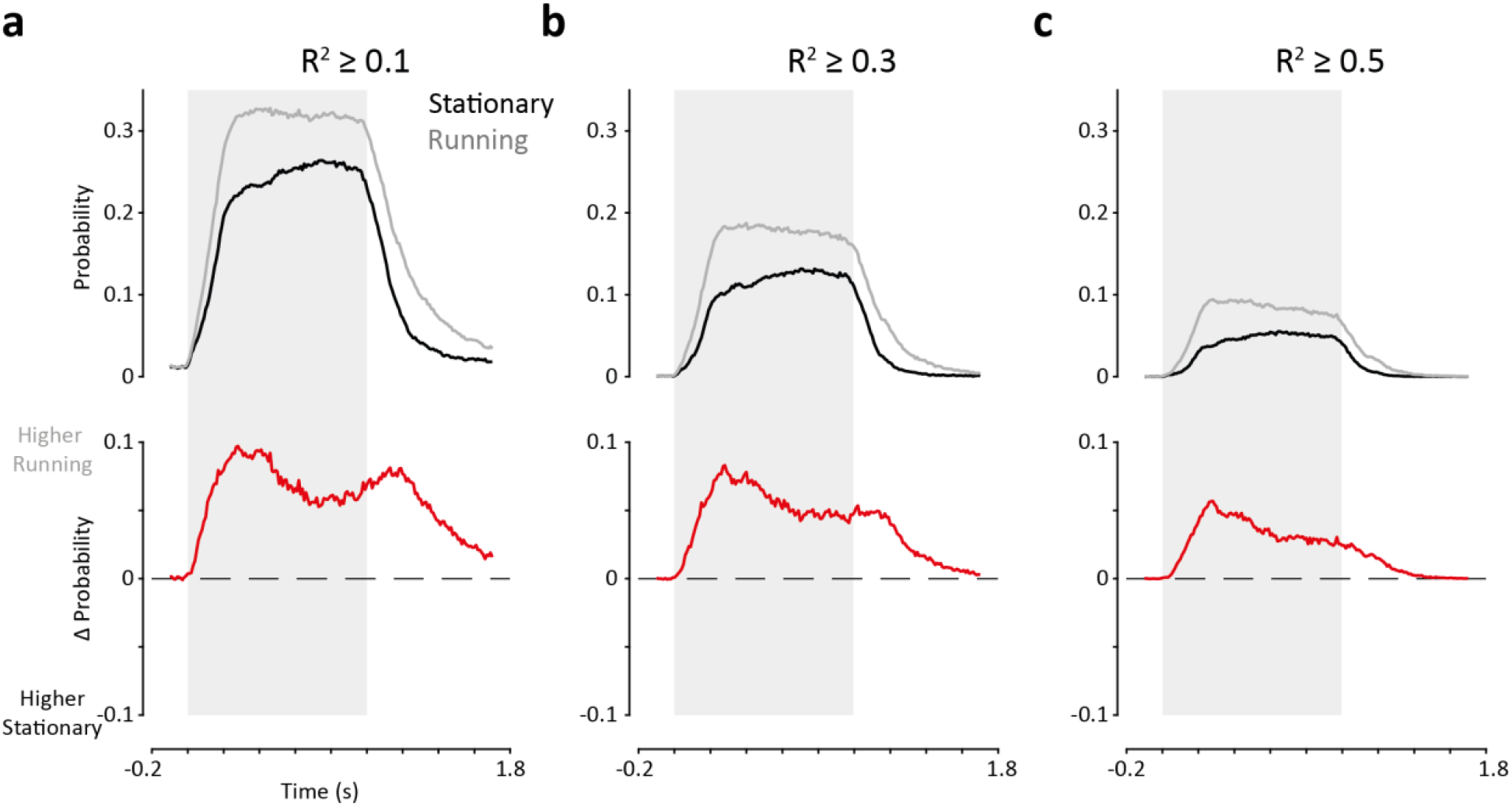
Fractions of neurons tuned for visual speed over time in stationary and running sessions. **a.** *Top panel:* Probability of a neuron (Stationary: n = 10,537; running n = 9,165) being tuned (R2≥0.1) as a function of time in stationary (black trace) and running (grey trace) sessions. Tuning strength was assessed using a sliding window (200ms window size, 10ms step size). *Bottom panel:* Difference in probability of a neuron being tuned between stationary and running sessions as a function of time (positive values indicate a higher probability in running sessions). Grey shaded region indicates stimulus presentation period. X-axis represents the window midpoint. **b-c.** Same as **a** for tuning strength thresholds of R2≥0.3 (**b**) and R2≥0.5 (**c**).

**Supplementary Figure 4.**
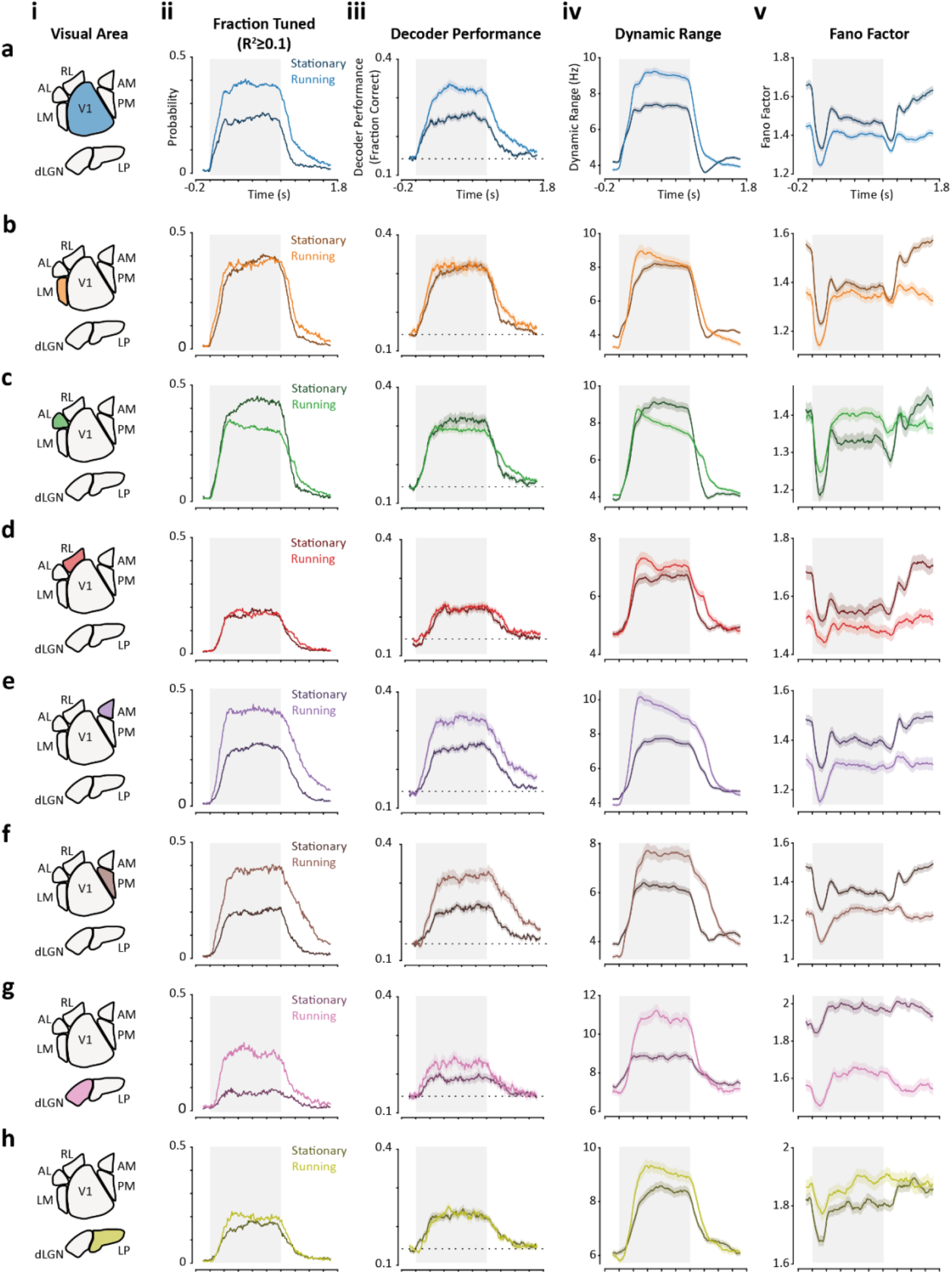
Fractions of tuned neurons, decoding performance, dynamic range and mean Fano factor over time in stationary and running sessions for each visual area. **a-h.** Visual speed encoding across time for neurons recorded in stationary and running sessions in each visual area.

i. Schematic of visual area anatomical location relative to other visual areas. Relevant visual area is colour-filled
ii. Related to **Supplementary Figure 3**. Probability of a neuron being tuned as a function of time for stationary (dark tone) and running (light tone) sessions (Number of sets of responses [stationary sessions / running sessions]: V1 [1,797 / 1,832], LM [1,760 / 854], AL [929, 2,012], RL [1,204 / 1,258], AM [1,812 / 1,174], PM [1,241 / 828], dLGN [620, 435], LP [1,174 / 772].
iii. Related to **Figure 2d**. Decoding performance (fraction correct responses) as a function of time. Dark and light shaded regions indicate mean ± sem performance of populations (n = 10 neurons) recorded in stationary and running sessions, respectively (Number of populations [stationary sessions / running sessions]: V1 [56 / 43], LM [51 / 24], AL [30 / 48], RL [36 / 35], AM [56 / 35], PM [33 / 19], dLGN [14 / 12], LP [30 / 20]). Dashed line indicates chance level performance.
iv. Related to **Figure 2e** (top panel). Dynamic range of single neuron responses (same sets of responses as [ii]) to the seven visual speeds presented as a function of time. Dark and light shaded traces indicate mean ± sem for neurons recorded in stationary and running sessions respectively.
v. Related to **Figure 2e** (bottom panel). Same as [iv] for the mean Fano factor of single neuron responses to the seven visual speeds presented.

**Supplementary Figure 5.**
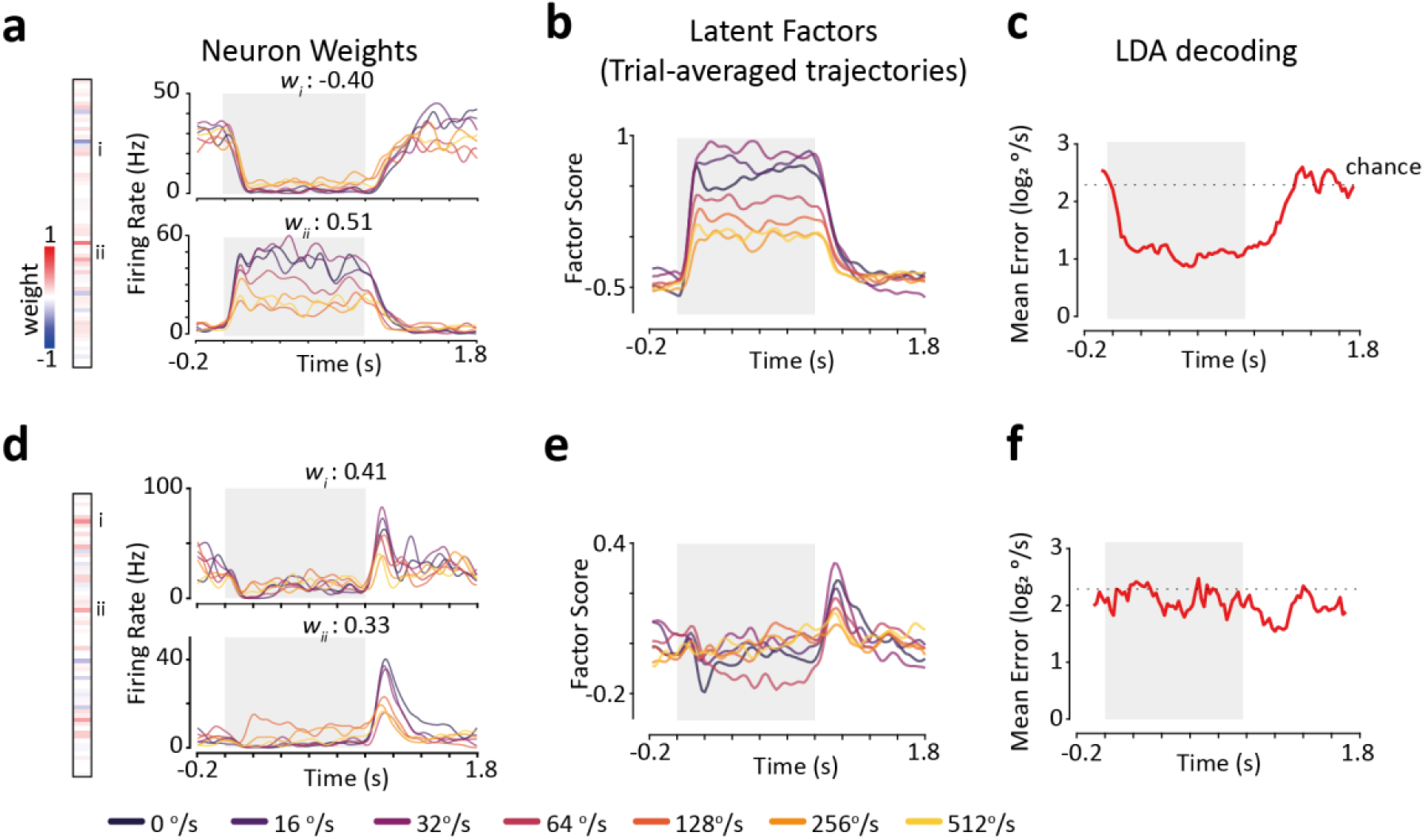
Example latent factors of population activity obtained using Factor Analysis. **a.** Illustration of an example latent factor. *Left panel:* factor loading pattern. A set of weights are assigned to neurons within a population for each latent factor. Each row represents the weight of an individual neuron. *Right panel*: PSTH responses to the seven visual speeds presented for two example neurons. The value above each plot represents the weight assigned to that neuron for the example latent factor. The relevant position on the factor loading pattern is also indicated. Grey shaded region indicates stimulus presentation period. **b.** Trial-averaged trajectories of the latent factor shown in **a** for each visual speed. **c.**Decoding performance over the response period using single-trial trajectories of the latent factor shown in **a** and **b**. Decoding was performed using LDA and a sliding window (200ms window size, 20ms step size). **d-f.** Same as **a-c** for a second example latent factor extracted from the same V1 population.

**Supplementary Figure 6.**
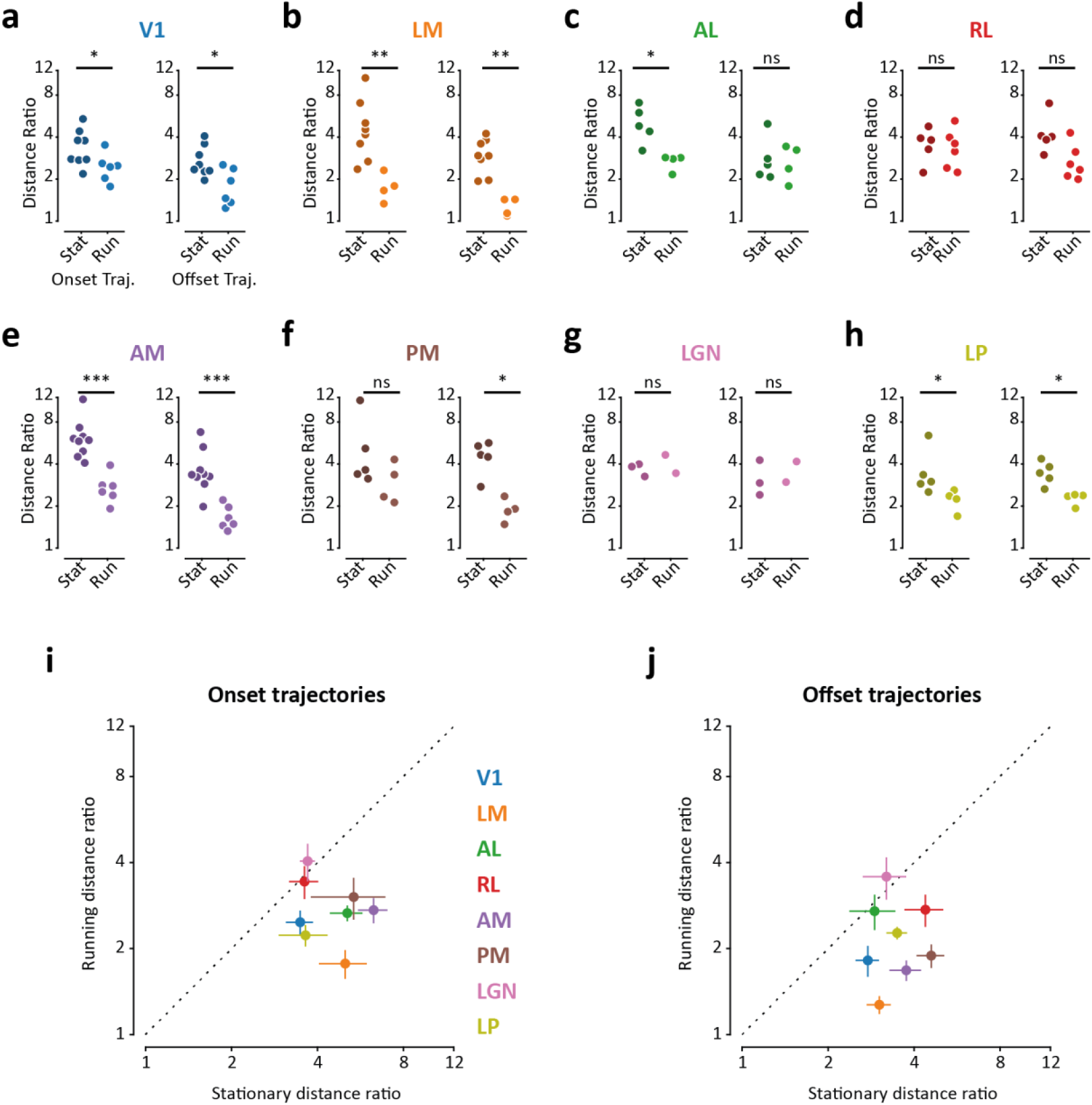
Latent trajectory Distance Ratios in stationary and running sessions for each visual area. **a-h.** Related to **Figure 3d, g** for each visual area. Mean Distance Ratio values for neural trajectories during the stimulus onset (*left panels*) and offset (*right panels*) periods, calculated for populations of neurons recorded in stationary (dark shade) and running (light shade) sessions Significance tested using Mann-Whitney U-test. **i.** comparison of mean Distance Ratio values (averaged over sessions) for the stimulus onset period for each visual area. **j.** same as **i** for stimulus offset. Errorbars are ± sem. *** p<0.001, ** p<0.01, * p<0.05, ns p≥0.05.

**Supplementary Figure 7.**
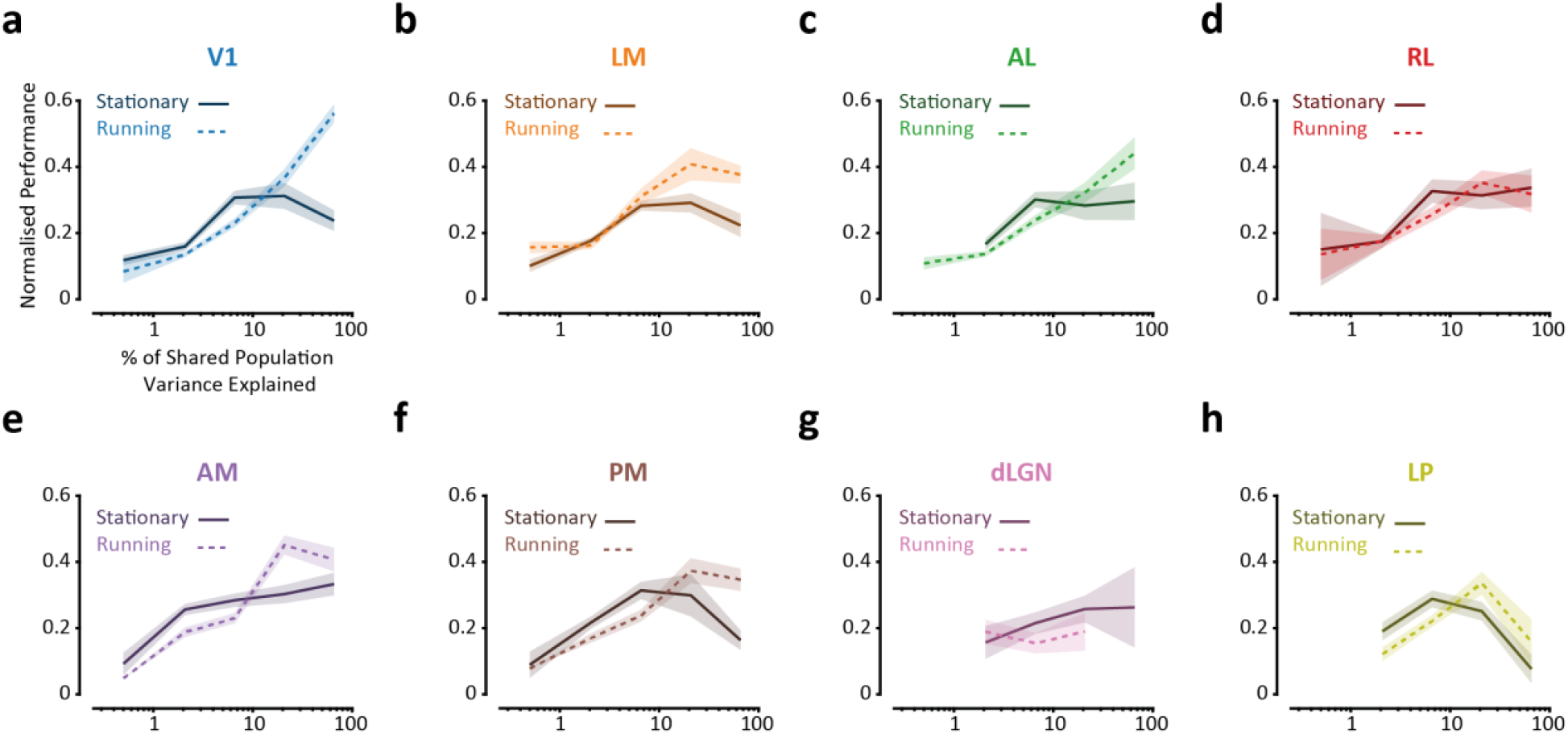
Relationship between decoding performance and % shared population variance explained in stationary and running states, for each visual area. **a-h.** Related to **Figure 5e** for each visual area. Mean normalised decoding performance of latent factors as a function of binned % shared population variance explained for populations recorded in stationary (dark shade) and running (light shade, dashed line) sessions (Number of populations [stationary / running]: V1 [22 / 24], LM [27 / 13], AL [13 / 16], RL [11 / 21], AM [27 / 21], PM [13 / 16], dLGN [8 / 4], LP [17 / 13]).

